# Forensic Facial Approximation of Achondroplastic Dwarf from Medieval Cemetery in Central Europe

**DOI:** 10.1101/2023.08.26.553833

**Authors:** Cicero Moraes, Marta Krenz-Niedbała, Sylwia Łukasik, Camilo Serrano Prada

**Author notes:** Correspondence to Cicero Moraes.

## Abstract

Achondroplasia (ACH, achondroplastic dwarfism) represents the most common form of skeletal dysplasia, occurring in c. 4 out of every 100,000 births. This study presents a computer-based facial approximation of the skull of a male individual suffering from ACH, who died at 30-45 years of age and was buried in Łekno, Poland between the 9th and 11th centuries AD. For the approximation procedure, soft tissue data from CT scans and ultrasonic measurements performed on living individuals were used. Additionally, the anatomical deformation technique was applied to arrive at the most reliable reconstruction of the dwarf’s appearance. To our knowledge, this is the first recreation of a person with achondroplasia, and one of the few showing a head of an individual suffering from a hereditary disease, with dimensions and shape differing from the population average values.

**Highlights:**

- Forensic facial approximation of an achondroplastic dwarf from 9th–11th century AD has been performed as the first in the world
- The applied procedure included CT of a virtual donor
- Anatomical deformation technique allowed to extract the endocast, revealing a large volume of the endocranium
- Few measurements applied to the facial skeleton proved successful in identification of a person suffering from achondroplasia

## 1. Introduction

### 1.1. Facial Approximation

Facial approximation, which is aimed to recreate a person’s facial appearance from the skull, is mainly used in three scientific disciplines: forensics, paleoanthropology and archaeology (Gietzen et al. 2019). In forensics it serves for personal identification, while in paleoanthropology and archaeology it is used to visualize the facial appearance of archaic humans or people from the distant past (Wilkinson 2010). The latter is mainly performed for educational and social purposes, such as dissemination of knowledge through reconstructions of famous historical figures, like Tutankhamen or King Richard III, and also to enable comparison of appearance of past and contemporary faces. There is also a growing number of reconstructions of common people from distant societies (https://www.livescience.com/gallery-of-reconstructions). Some applications of facial approximation in anthropological and medical sciences refer to the interpretation of trauma and disease, in which they constitute an aid in the analysis of facial appearance relating to ancient diseases, medical treatment, and hereditary conditions (Wilkinson 2010), such as in the case of facial trauma of Philip II Macedon from ancient Greece, and a medieval English soldier exhibiting a healed injury to the mandible (Wilkinson, Neave 2003).

The reconstruction techniques can be two dimensional (2D) or three dimensional (3D), and either manual or computerized (Gupta et al 2015). With the advancement in 3D and software technologies computer-based methods have been increasingly applied, because they can provide consistent and objective, efficient, and cost-effective results (Gietzen et al. 2019). This approach brings together anthropology, osteology, anatomy, computer science and archaeology not only to obtain the most accurate anatomical reconstruction, but also to take into account such features as hairstyle, pigmentation, and clothing (Wilkinson 2010).

### 1.2. Achondroplasia

Achondroplasia (ACH, achondroplastic dwarfism) represents the most common form of skeletal dysplasias, which is also the most frequently found type of dysplasias in bioarchaeological record. It belongs specifically to chondrodysplasias, which result from genetically determined abnormal cartilage formation (Lewis 2019). Achondroplasia is a congenital, hereditary, and familial disease with an autosomal dominant mode of inheritance, but is most commonly caused by a de novo germ cell mutation (Foreman et al. 2020). Worldwide there are about 250,000 affected people (Pauli 2019). In modern European populations the pooled birth prevalence per 100,000 median is 3.5 (Foreman et al. 2020), and specifically in Poland, Wielkopolska region, the area the individual discussed in this study originates from, it is 4.47 per 100,000 births (Coi et al. 2019). The most obvious result of severe inhibition of cartilage proliferation in achondroplasia is reduced body height, which is 130 cm on average in adult males and 125 cm in females (Pauli 2019). It should be emphasized that the stature of males in the medieval population of Łekno involved in this study was on average 175.8 cm, based on the formula for femur bicondylar length (Vercellotti et al. 2009). Other clinical features include disproportional shortening of the limbs relative to the trunk (rhizomelic disproportion), thoracolumbar kyphosis, lumbar hyperlordosis, joint hypermobility, but limited extension and rotation of the elbow and hip, short fingers and so called “trident” hand, with shortened fingers and tapering terminal phalanges. The skull is typically large (macrocephalic) and robust, with frontal bossing, depressed nasal bridge and midfacial area. The majority of affected persons have rather normal life expectancy, although in mid-adulthood there is an increased risk for premature death due to cardiovascular complications (Pauli 2019, Julie Hoover-Fong et al. 2021).

The aims of this study were to apply the forensic facial approximation method, based on the head model of a healthy living person, to reconstruct the appearance of an achondroplastic dwarf from a medieval Central European population, to compare chosen cranial metric characteristics of the examined dwarf with the data on healthy individuals, and to apply a method of calculating endocranial volume to discuss this dimension in the achondroplastic vs healthy male.

## 2. Material and methods

### 2.1. Examined individual

The present study used two 3D models of the skull belonging to the individual Ł3/66/90 recovered in 1990 from a medieval cemetery in Łekno, Poland, and dated to the 9th–11th century AD. The excavated skeletal remains represent a male individual with a stature of 115 cm, who died at 30-45 years of age (Miłosz 1993; Matczak et al. 2022). Research performed by Matczak et al. (2022) revealed that he suffering from achondroplasia, Leri-Weill dyschondrosteosis and ulnar hemimelia.

### 2.2. 3D scanning and model processing

The examined skull was scanned using a high-resolution Artec Space Spider 3D scanner based on blue-light technology. Bones were gently rotated 360 degrees on a turntable and scanned in real-time from different angles collecting data covering the area of the whole skull, except two loose teeth (FDI 32, 42), which were not scanned. Applied technology allowed for capturing high-resolution geometry and texture of scanned bones with a 3D resolution of up to 0.1 mm and 3D point accuracy of up to 0.05 mm (https://cdn.artec3d.com/pdf/Artec3D-SpaceSpider.pdf). To generate 3D models Artec Studio 15 Professional software was used (Artec 3D, Luxembourg). Processing 3D scans was performed manually. Applying the “Don’t fill” option in the hole-filling method allowed for leaving the natural anatomical openings in the skull. Then, prepared models were exported to .ply extension and uploaded to the Sketchfab platform as supporting information for research published elsewhere (see Matczak et al. 2022). Both created models are available on the Sketchfab account of the AMU Human Evolutionary Biology Research Team (https://sketchfab.com/lukasik) under the names: Achondroplastic dwarf - Skull (https://skfb.ly/oB6qv) and Achondroplastic dwarf - mandible (https://skfb.ly/oB9Wu).

### 2.3. Anatomical analysis of 3D skull model

The human skull presents as follows: brachycephalic aspect, without bone fissures in the massive facial skull, without irregularities on the bone surface, showing a frontal prominence marked by the absence of a nasal bridge, symmetrical orbits with an anti-mongoloid arrangement, hypoplasia in the middle third of the face with accentuated concavity in the bilateral maxillomalar pillar, nasal septum with deviation to the left in its caudal portion, pyriform aperture and choanae without stenosis, emergence of the supraciliary, infraorbital and mental nerves, in addition to superior and inferior orbital fissures with bilateral nasolacrimal ducts present. Maxillomandibular relationship with mandibular prognathism, hypergonia and what appears to be a fracture line in the mandibular symphysis. Appearance of atypical condylar morphology, finding flat condylar surfaces and glenoid fossa, bilateral external auditory canal present. Upper teeth with occlusal surfaces without recognizable anatomy, absent (FDI) 11,12,13,21,18 and 28,31,32,36,41,42,46. A cavitation lesion can be seen on teeth 26 and 37 with coronal fracture on tooth 43.

### 2.4. Facial reconstruction procedure

The present work uses the same step-by-step approach discussed in Abdullah et al. (2022), starting with the complementation of the missing regions of the skull (see Anatomical analysis of 3D skull model), followed by the projection of the profile and structures of the face from statistical data (Moraes et al. 2021, Moraes et al. 2022c, Moraes & Suharschi 2022) generating the volume of the face with the aid of the technique of deformation/anatomical adaptation and the finishing with the detailing of the face, configuration of the hair and generation of the final images.

All the facial approximation work was carried out in the Blender 3D software (www.blender.org), using the add-on OrtogOnBlender (http://www.ciceromoraes.com.br/doc/pt_br/OrtogOnBlender/index.html), which expands the potential of the native software, such as the possibility of importing and reconstructing computed tomography scans. The add-on has a submodule specialized in facial approximation, called ForensicOnBlender (https://github.com/cogitas3d/OrtogOnBlender/blob/master/ForensicOnBlender.py). Both Blender and the OrtogOnBlender add-on are open source and free, and run on the most popular operating systems currently available: Windows, Linux and Mac OSX.

The skull aligned to the Frankfurt horizontal plane received a series of soft tissue thickness markers, corresponding to 31 anatomical points, measured in living individuals with ultrasound (De Greef et al. 2006). Roughly speaking, these markers limit the skin in a series of regions, but do not cover some such as the nose, for example (Figure 1A). Three approaches were used to project the nose by Prokopec and Ubelaker (2002), by Gerasimov (1971) (Stephan et al 2003) and by Moraes et al. (2021) based on lifting the nose in relation to the bones, through computed tomography. There was a very large discrepancy (Figure 1B), between the projections for 1971 and 2002 in relation to that for 2021. As this is an individual with a face outside the pattern studied, initially it was decided to draw a nose in the average of the projections, but even so it was left out of the standard deviation of the newest study, based on CT scans (Figure 1B). To complement the coherence of the approximate structure, the reconstructed computed tomography of a virtual donor (Brazilian with European ancestry) was imported containing the 3D meshes of soft tissue and bones. This structure was deformed in order to make the donor’s skull compatible with the Ł3/66/90 skull. When making the deformation/adaptation in the bones, this restructuring reflects in the soft tissue, generating an approximate and anatomically compatible face with a real head (Figure 1 C-E), in order to complement the projections made with the soft tissue markers and nose, which despite being very coherent, do not cover all the structures to be approximated. When observing the projection of the nose with the anatomical deformation, it is attested that the first was smaller than the second. The anatomical deformation allows adjusting the outline of the nose and chin (Figure 1F). When observing the nose tracing more carefully, it is noticed that the adjustment is compatible with the projection from tomography data (Moraes et al 2021), coinciding with the standard deviation limit (Figure 1G), this may indicate that, even when dealing with a face with part of the structure different from the general average, the proportion of the nose respects the expected projection. The same cannot be said about the foramen magnum, which resulted in a significantly smaller structure (Hecht et al 1985) than that of the virtual donor (Figure 1C). To assess the structure of the soft tissue, some anatomical points were placed on the skull, so that a series of lines corresponding to the limits of the lips, ears, position of the eyeballs, nasal wings, lower limit of the nose, size of the eyes and others. Such projections come from studies carried out with CT scans of living individuals (Moraes et al. 2022a, Moraes et al. 2022c, Moraes & Suharschi 2022) and the result in the approximation of the individual Ł3/66/90 was within the expected parameters (Figure 1H). Following the approach proposed in the studies by Abdullah et al. (2022) (Abdullah et al. 2022), a mesh from another facial approximation was used and deformed/adapted to be compatible with the current work, not only in the shape of the face (Figure 1I), but in the configuration of the beard and hair (Figure 1J).

**Figure 1.**
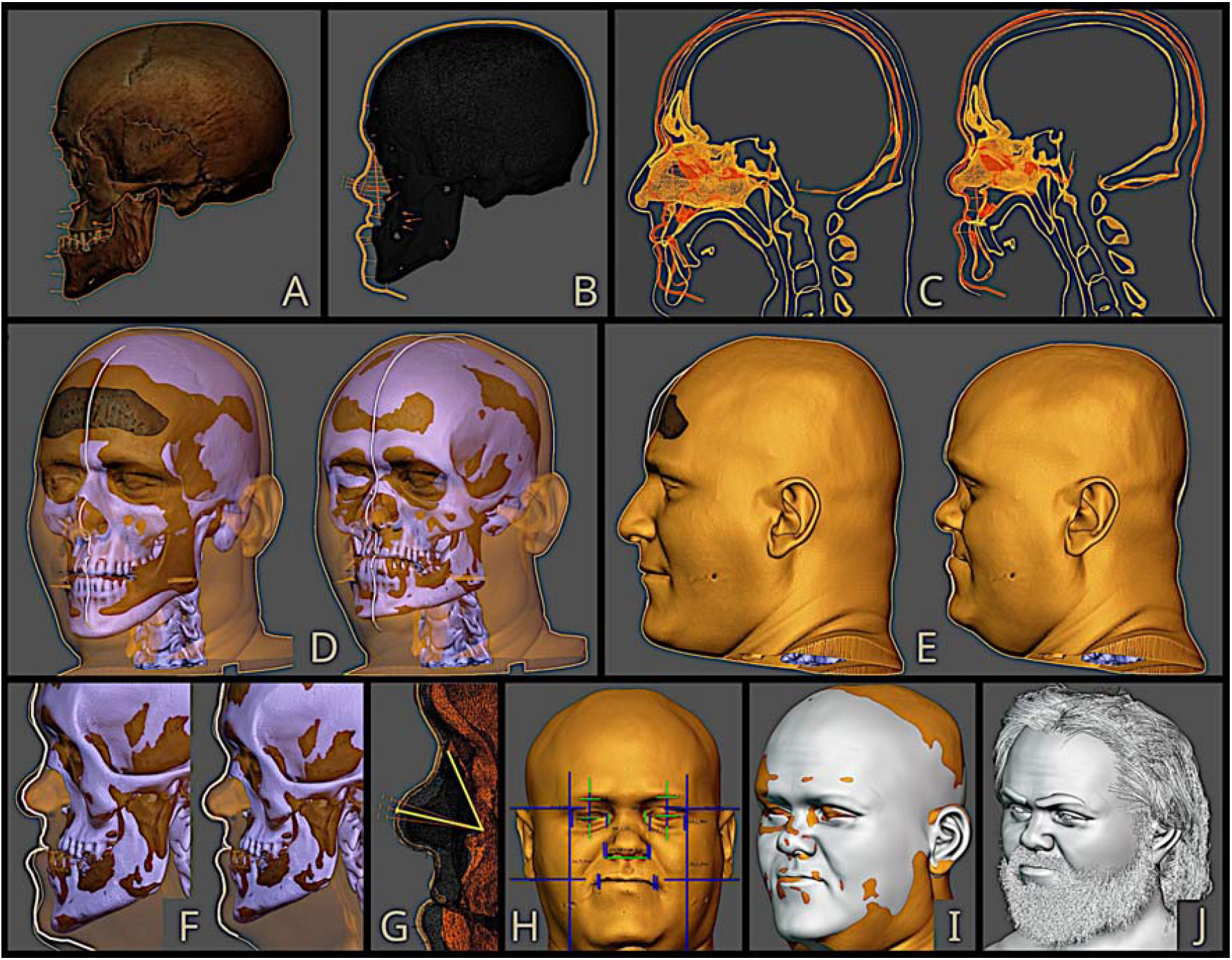
Steps of the facial approximation of the examined individual: A) Skull with soft tissue thickness markers, B) Skull with the projected nose and face profile, C) Head CT of the virtual donor over Ł3/66/90 skull (left), and after the anatomical deformation (right), D) Approximation face of the dwarf (right) from the virtual donor (left), E) face reconstruction of the virtual donor (left) and the dwarf (right), F) Interpolation of face profile line from the projected line (left) and using anatomical deformation (right), G) The interpolated profile line is compatible with the nasal projection from statistical data, H) Face measurements performed, I) Face with preconfigured texture and hair, J) Reconstructed face of the dwarf with hair and beard.

The endocast of the Ł3/66/90 skull was segmented with the 3D mesh editing tools of the Blender software, the volume of the resulting mesh was extracted from the add-on 3D Print Toolbox (https://docs.blender.org/manual/en/latest/addons/mesh/3d_print_toolbox.html) and the other measurements were performed with the Measureit add-on (https://docs.blender.org/manual/en/latest/addons/3d_view/measureit.html).

## 3. Results

### 3.1. Dwarf Facial Approximation

In the first step of the applied facial approximation method using a deformation/adaptation approach based on a real donor’s head, a dwarf’s head appearance has been recreated without specific individual hair features in order to show the details of the face, in particular the mandibular prognathism (class III). A maxillary relationship is identified as skeletal class III anomaly due to the marked hypoplasia of the midface with maxillary retroposition respect to the mandible, constituting a marked concave facial profile, this jaws disposition and the marked maxillary hypoplasia reflects a severe dental class III malocclusion too with deep bite. It is noteworthy that the mandibular condyles in the specimen are incomplete, so it is possible that the dentofacial anomaly in life was more severe than what is seen in the reconstruction (Jacobson et al. 1974). We generated an image based on a more simplified color palette that resembles a sepia tone (Figure 2).

**Figure 2.**
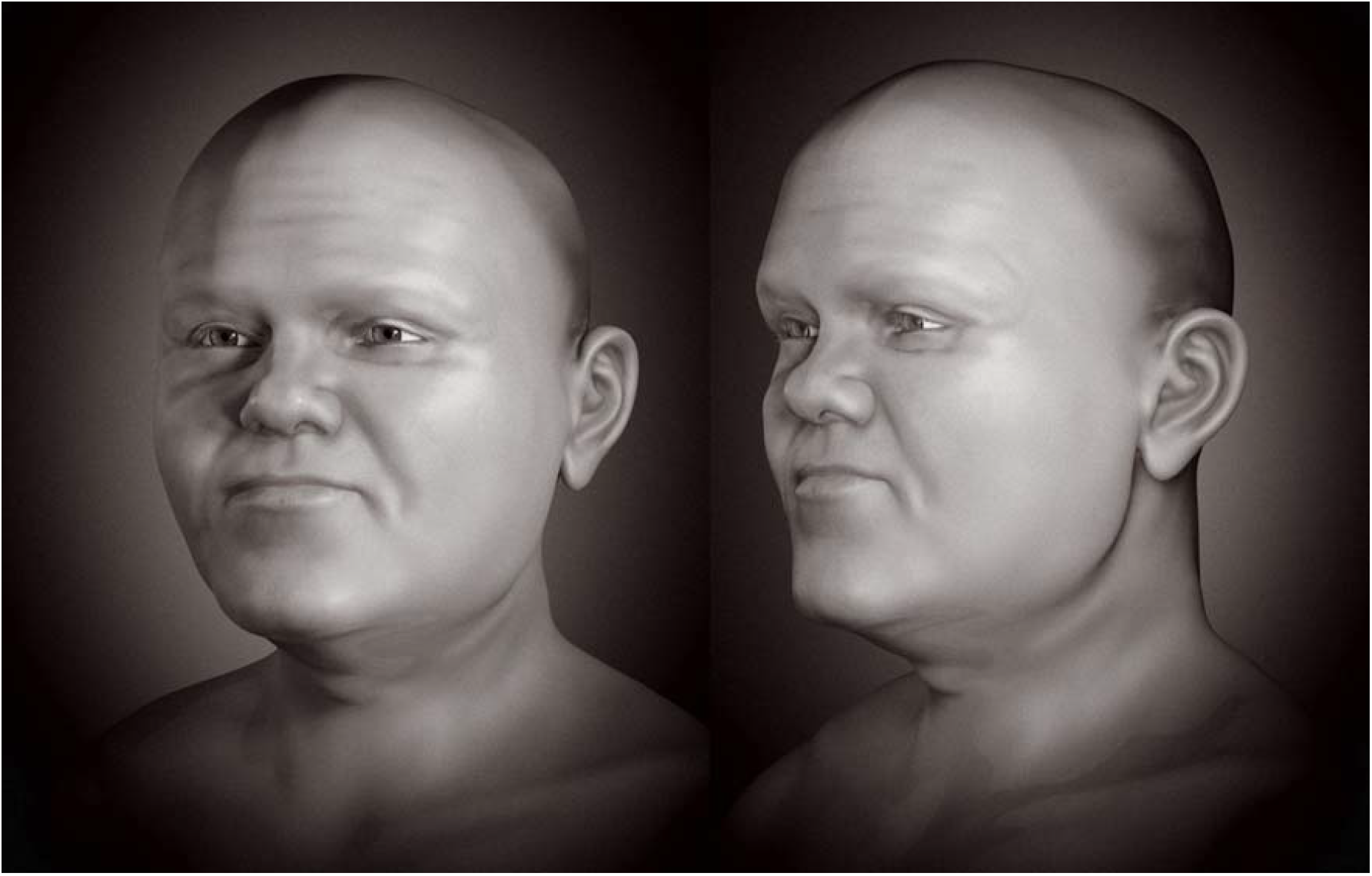
Face of the individual examined without hair and beard.

In the second step hair features have been added. The image with beard, eyebrows and hair, even though it is more subjective, humanizes the individual, allowing better recognition by the general population, not specialists in forensic sciences or even archeology (Figure 3).

**Figure 3.**
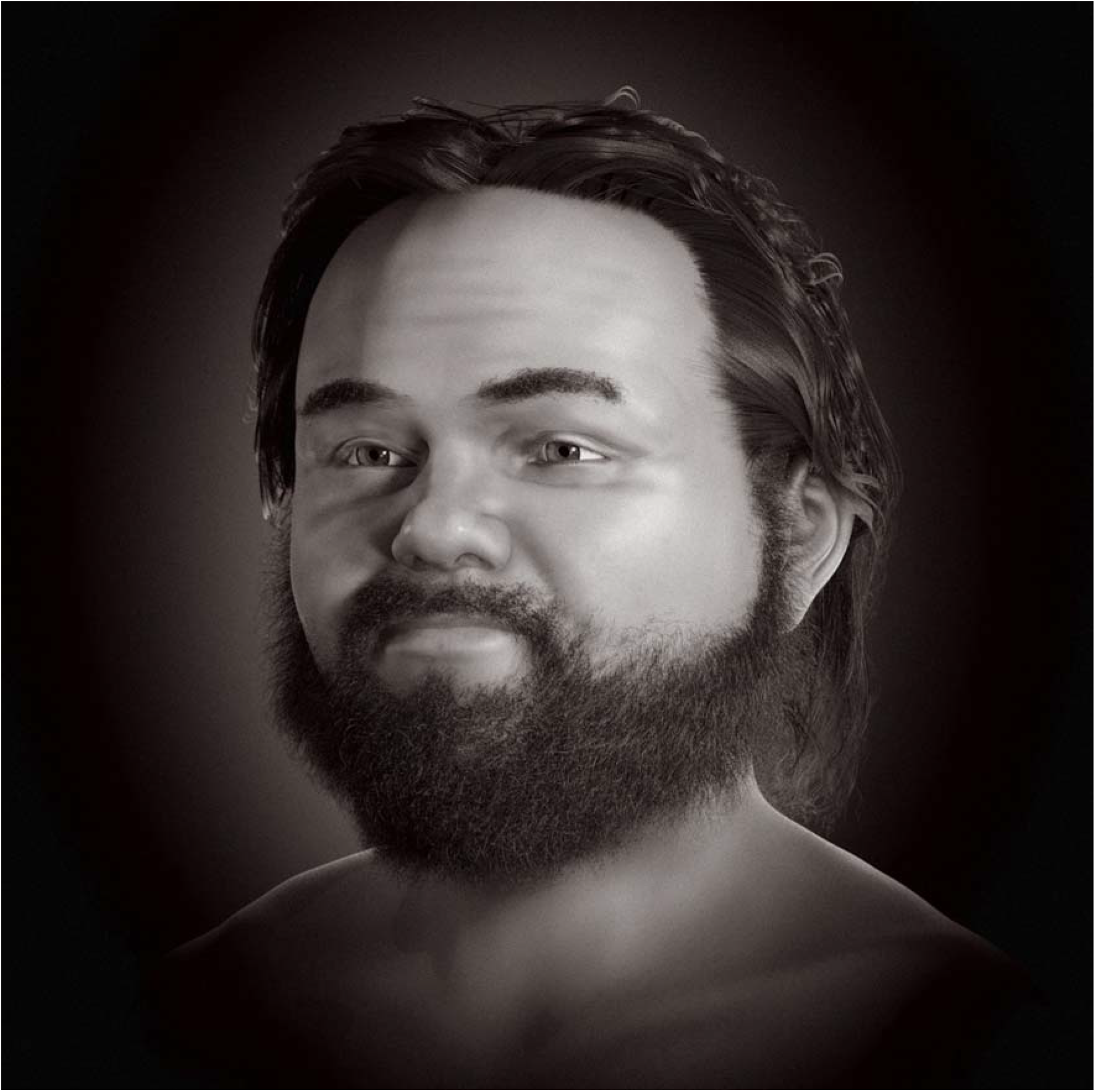
Face of the individual examined with hair and beard.

### 3.2. Skull dimensions of dwarf and individuals unaffected by ACH

The authors took advantage of the opportunity to analyze the population distribution factor based on proportions based on the frontomalar orbital distance (fmo-fmo) (Moraes et al. 2022d). The sample of 213 skulls includes the ACH case analyzed in the present study (Figure 4). It was possible to attest that the examined individual was positioned in a separate location against unaffected people. More studies are needed to attest to such a trend, involving other cases of achondroplasia to take into account a greater range of biological variation.

**Figure 4.**
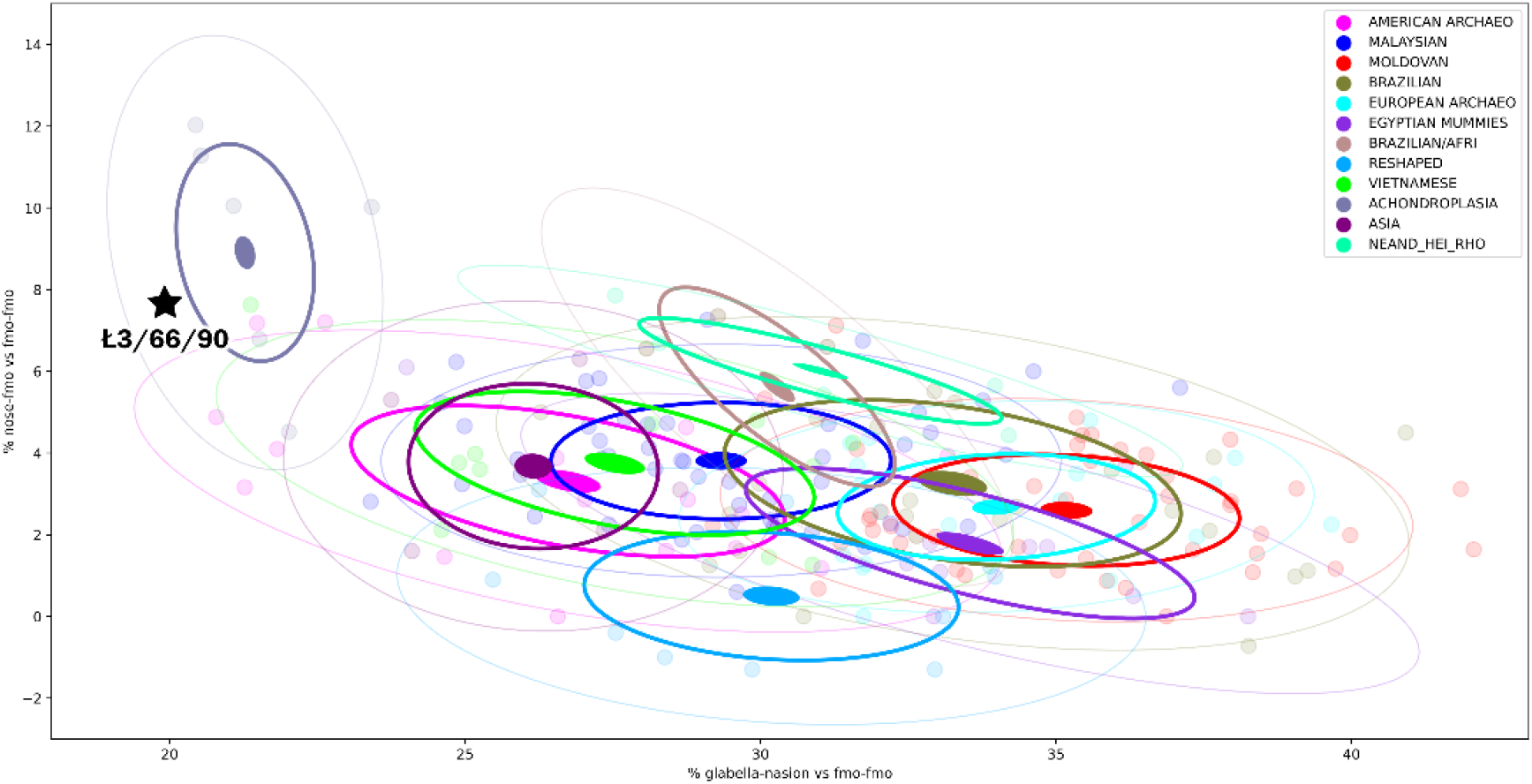
Positioning of the Ł3/66/90 skull against unaffected individuals. The X axis are the value of the factor ((g-n)*100)/fmo-fmo), and the Y axis are the values of the factor ((rhi-ec)*100)/fmo-fmo)(source of comparative data Moraes et al. 2022d).

### 3.3. Endocranial volume

Once the virtual donor was deformed/adjusted to the Ł3/66/90 skull, it was possible to extract the endocranium (Abdullah et al. 2021) (Figure 5) and measure the volume at 1676 ml, which turned out to be two standard deviations above the mean of 1328 (±164 ml) in a study carried out with modern human skulls (Neubauer et al. 2018). The approximate soft-tissue head circumference (60.99 cm) was also two standard deviations above the expected mean for adult males (≈57 ± 1 cm) (PAHO, 2020).

**Figure 5.**
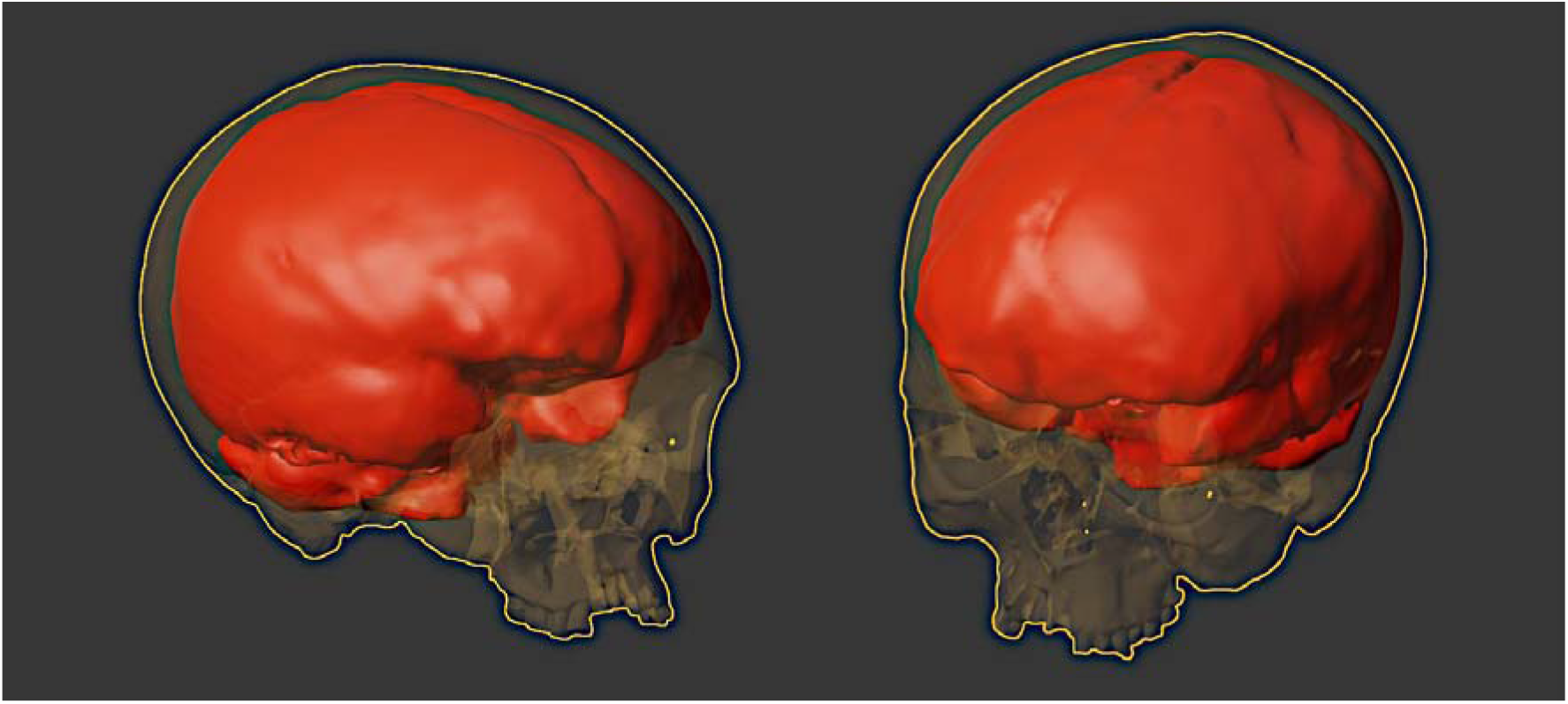
Visualization of the endocranium of the Ł3/66/90 individual.

## 4. Discussion

To our best knowledge, there are not any other facial approximation of an individual with achondroplasia, which would situate this study as the first of this kind in the world. With our work we would like to emphasize, that even in cases of serious skeletal deformations facial appearance can be effectively recreated.

Facial approximation techniques are based on population averages, which are composed mostly of individuals of normal height. Thus, because we deal here with a skeletal dysplasia case the approach applied in our paper could be treated with caution. However, nasal projection techniques by proportion of distances and anatomical deformation circumvent the limitations of conventional methods of facial approximation Russian, Manchester and American. They are subject to interpretation of particular anthropologists and subjective experience of the person performing the manual work. As such multiple reconstructions of the same skull can be produced (Shui et al. 2021). In contrast our computerized approximation is fully based on statistical data, extracted from living people. More studies are needed to confirm the accuracy of the method, which may be difficult to implement given the rarity of individuals with the same Ł3/66/90 skull pattern.

By publicizing works such as the present one, there is a possibility of putting conditions such as achondroplasia up for public debate, which highlights the facial approximation as a popular communication tool, aiming at getting the public interest. The facial approximation itself benefits from such a project, since the work does not focus only on known historical figures, but on anonymous individuals with specific conditions such as dwarfism, syphilis (Jackson, 2022), cranial remodeling (Solly, 2023), and others.

In terms of performed cranial measurements, the examined individual with achondroplasia took a separate position in the plot. Thus it seems that the measures and factors used are good parameters for distinguishing people with such condition, which allows researchers to make such a projection even if they do not have a complete skull available. The most obvious limitation for such an approach is that it is based on only one individual with achondroplasia. More studies are needed to account for potential greater biological variation, so that a more robust scenario can be formed.

## 5. Conclusions

Our research for the first time showed a recreation of the facial appearance of an achondroplastic dwarf based on 3D models of the skeletal remains and using the CT image of a virtual donor. This is an average image of an individual suffering from ACH performed with the use of European average soft tissue measurements. We found that the cranial metric characteristics of bioarchaeological remains of ACH individual are markedly different from healthy persons. Intracranial volume of the dwarf exceeds two standard deviations above the mean of unaffected modern people.

## Acknowledgments

To Dr. Richard Gravalos (Consultório Dr. Richard Gravalos, São Paulo, Brasil) for providing the CT scan of the virtual donor, used in the anatomical deformation/adaptation.

## Author contributions

Conceptualization: C.M., M.K.N., S.Ł.; data curation, S.Ł., M.K.N, methodology and visualisation: C.M., writing—original draft, C.M., M.K.N., S.Ł., C.S.P.; writing— review and editing, C.M., M.K.N., S.Ł.

## Conflict of interests

The authors declare no conflict of interest.

## Data statement

The data that support the facial approximation computerized procedure used in this study are available from the corresponding author upon reasonable request. 3D models are uploaded to the Sketchfab platform https://sketchfab.com/lukasik/collections/achondroplastic-dwarf-eabdf52c8b2d4cabb63d0b9cf1f374e2.

## Notes

### Competing Interest Statement

The authors have declared no competing interest.

